# Precision engineering of an anti-HLA-A2 chimeric antigen receptor in regulatory T cells for transplant immune tolerance

**DOI:** 10.1101/2021.08.16.456548

**Authors:** Yannick D. Muller, Leonardo M.R. Ferreira, Emilie Ronin, Patrick Ho, Vinh Nguyen, Gaetano Faleo, Yu Zhou, Karim Lee, Kevin K. Leung, Nikolaos Skartsis, Anupurna M. Kaul, Arend Mulder, Frans H.J. Claas, James A. Wells, Jeffrey A. Bluestone, Qizhi Tang

## Abstract

Infusion of regulatory T cells (Tregs) engineered with a chimeric antigen receptor (CAR) targeting donor-derived human leukocyte antigen (HLA) is a promising strategy to promote transplant tolerance. Here, we describe an anti-HLA-A2 CAR (A2-CAR) generated by grafting the complementarity-determining regions (CDRs) of a human monoclonal anti-HLA-A2 antibody into the framework regions of the Herceptin 4D5 single-chain variable fragment and fusing it with a CD28-ζ signaling domain. The CDR-grafted A2-CAR maintained the specificity of the original antibody. We then generated HLA-A2 mono-specific human CAR Tregs either by deleting the endogenous T-cell receptor (TCR) via CRISPR/Cas9 and introducing the A2-CAR using lentiviral transduction or by directly integrating the CAR construct into the TCR alpha constant locus using homology-directed repair. These A2-CAR^+^TCR^deficient^ human Tregs maintained both Treg phenotype and function *in vitro*. Moreover, they selectively accumulated in HLA-A2-expressing islets transplanted from either HLA-A2 transgenic mice or deceased human donors. A2-CAR^+^TCR^deficient^ Tregs did not impair the function of these HLA-A2^+^ islets, whereas similarly engineered A2-CAR^+^TCR^deficient^CD4^+^ conventional T cells rejected the islets in less than 2 weeks. A2-CAR^+^TCR^deficient^ Tregs delayed graft-versus-host disease only in the presence of HLA-A2, expressed either by co-transferred peripheral blood mononuclear cells or by the recipient mice. Altogether, we demonstrate that genome-engineered mono-antigen-specific A2-CAR Tregs localize to HLA-A2-expressing grafts and exhibit antigen-dependent *in vivo* suppression, independent of TCR expression. These approaches may be applied towards developing precision Treg cell therapies for transplant tolerance.

## Introduction

Regulatory T cells (Tregs) are a small subset of CD4^+^ T cells that are key for maintaining self-tolerance and preventing autoimmune disease (1). A plethora of preclinical models have shown that the infusion of Tregs can suppress graft rejection and promote transplant tolerance (2). Several phase I/II clinical studies using Tregs have been reported (3, 4). The ONE Study is the largest coordinated international study of regulatory cell therapies in kidney transplantation. The study includes 28 patients who received Treg therapy in 4 non-randomized single-arm phase I/IIa trials. The results demonstrated feasibility, safety, and potential benefit of Treg-based therapies to reduce the burden of immunosuppression (5). While a significant fraction of Tregs in the polyclonal pool can react to allogeneic donor antigens, data from preclinical models show that donor-reactive Tregs are more effective than polyclonal Tregs in promoting transplant tolerance (6). Unfortunately, donor alloantigen-reactive Tregs may be functionally altered or induced to migrate out of the peripheral blood following transplantation, thus limiting the frequency of alloantigen-reactive clones within polyclonal Treg products and thereby posing challenges for consistent expansion of donor-reactive Tregs (2).

Redirecting Treg alloantigen reactivity using an alloantigen-specific chimeric antigen receptor (CAR) may present a more reliable approach to enhancing therapeutic Treg donor reactivity and potency (7–9). Previous studies have shown that a CAR, consisting of a mouse anti-HLA-A2 (A2) antibody-derived single chain variable fragment (scFv) coupled to a CD28-ζ signaling domain, could be introduced in human Tregs using lentivirus. These A2-CAR Tregs demonstrated superior efficacy in preventing xenogeneic graft-versus-host disease (GvHD) in NSG mice when compared to polyclonal Tregs or Tregs transduced with an irrelevant CAR (10). The therapeutic potential of A2-CAR Tregs for organ transplantation was subsequently demonstrated by two separate groups which independently applied A2-CAR Tregs to prevent the rejection of A2^+^ human skin grafts in humanized mouse models, further bolstering the enthusiasm for evaluating this technology in humans (11, 12).

In all these studies, the CAR constructs were introduced into the Tregs via lentivirus and the engineered Tregs also expressed their endogenous TCR. Lentiviral transduction results in random integration of the CAR construct in the genome that can lead to variable levels of CAR expression, transcriptional silencing, or accidental disruption of important genes. A previous study has shown that site-specific integration of a CD19-CAR into the TCR alpha constant region (*TRAC*) of T cells results in a more uniform distribution and TCR-like regulation of CAR surface expression, thereby mitigating T-cell exhaustion and enhancing anti-tumor activity (13). In addition, we recently observed that CAR^hi^ human T cells exhibited a surprisingly robust proliferative response to anti-CD28 stimulation alone, independent of CAR or TCR engagement, whereas CAR^lo^ T cells did not (14). Thus, lentivirally engineered Tregs may result in heterogeneous CAR expression and unexpected properties of the engineered cells. Knocking a CAR into the *TRAC* locus and deleting the endogenous TCR may more precisely control CAR Treg activity. However, it is unclear whether CAR Tregs can function without the endogenous TCR. We thus conducted the current study by generating TCR^deficient^ A2-CAR human Tregs and assessed their trafficking, survival, and function in humanized NSG mouse hosts.

## Materials and Methods

### Human peripheral blood products and T cell isolation and expansion

Human peripheral blood from de-identified healthy donors was purchased from STEMCELL Technologies (Vancouver, Canada), which collects and distributes de-identified human blood products with consent forms, according to protocols approved by the Institutional Review Board (IRB). Peripheral blood mononuclear cells (PBMCs) were isolated by Ficoll (GE Healthcare, Chicago, IL) density gradient centrifugation. T cells were further enriched using the EasySep Human T Cell Isolation Kit (STEMCELL Technologies), as per the manufacturer’s instructions. Enriched CD3^+^ T cells, or CD4^+^CD127^+^CD25^low^ conventional T cells (Tconv) or CD4^+^CD127^low^CD25^high^ regulatory T cells (Tregs) purified by fluorescence-assisted cell sorting (FACS) using a BD FACS Aria II Cell Sorter (Beckton Dickinson, Franklin Lakes, NJ) were used for experiments. Tregs were expanded as previously described (15). Antibodies utilized for flow cytometry are summarized in Supplementary Table 1.

### Cloning and specificity verification of an anti-HLA-A2 scFv

A human B-cell derived hybridoma (clone SN607D8) was used as source material to produce an anti-HLA-A2 scFv. This hybridoma produces an IgG1κ monoclonal antibody that recognizes HLA serotypes A2 and A28 (16). RNA from the SN607D8 hybridoma was used as template for RT-PCR amplification of the V_L_ and V_H_ chains of the IgG. The scFv gene was then constructed in a V_H_-(GGGS)_3_linker-V_L_ format and incorporated into the pHEN1 phage display vector (17). The binding activity of phage-displayed scFv was assessed using two tumor cell lines, THP-1 (HLA-A*02:01/02:01, HLA-B*15:11/15:11 (18)) and RPMI 8226 (HLA-A*30:01/68:02, HLA-B*15:03/15:10 (19)). Binding to these cell lines was measured using sequential staining with a biotinylated anti-phage antibody and fluorochrome-conjugated streptavidin followed by flow cytometric analysis.

### Grafting of the anti-HLA-A2 scFv

The CDR regions of the anti-HLA-A2 scFv from hybridoma SN607D8 were grafted onto the 4D5 human antibody scaffold used in herceptin (trastuzumab) by pairwise alignment of amino acid residues using the software Jalview (20). The specific CDR3 regions of the anti-HLA-A2 scFv were predicted using the software Paratome (21). The grafted scFv was constructed in the V_H_-(GGGGS)_3_linker-V_L_ format.

### Lentivirus production

The A2-specific CAR was created by generating a chimeric DNA sequence encoding a MYC-tag upstream of the grafted anti-HLA-A2 scFv, an IgG4 hinge, CD28 transmembrane domain, and a CD28-CD3zeta tandem signaling domain (purchased as gblocks from Integrated DNA Technologies, IDT, Coralville, IA). The resulting DNA fragment was subcloned into a pCDH lentiviral vector containing an EF1α promoter (addgene-plasmid-64874 (22)). The CAR construct was linked to a truncated EGFR (EGFRt) or a luciferase gene via a 2A self-cleaving peptide sequence. All constructs used in subsequent experiments were confirmed by Sanger sequencing. Lentivirus was produced as previously described (23). Briefly, HEK293T cells were seeded at 3 × 10^6^ cells on 10 cm cell culture dishes 24 hours prior to transfection with 4 μg of plasmid DNA, 2 μg of the packaging vector pCMV-dR8.9, 2 μg of VSV envelope vector pMD2.G and 15 nmol linear 25 kDa polyethylenimine (Millipore Sigma, Burlington, MA). Media was replaced 24 hours later and ViralBoost Reagent (Alstem, Richmond, CA) was added. The supernatant was collected 24 and 48 hours later. Virus was concentrated using LentiX concentrator (Takara, Shiga, Japan).

### AAV6 production

A pAAV-MCS plasmid containing inverted terminal repeats (ITRs) from AAV serotype 2 (Agilent Technologies, Santa Clara, CA) was utilized as backbone for AAV6 plasmid construction (naturally occurring AAV6 has an AAV2 ITR (24)). Cloning was performed with in-fusion cloning tools and protocols provided by Takara. Large scale DNA preparation was performed using a Zymopure plasmid maxiprep kit (Zymo Research, Irvine, CA). All constructs used in subsequent experiments were confirmed by Sanger sequencing. For AAV production, 30 μg of pDGM6 helper plasmid (a gift from Dr. YY Chen, University of California, Los Angeles), 40 μg of pAAV helper, and 15 nmol linear polyethylenimine were used. AAV6 vector production was carried out by iodixanol gradient purification (25). After ultracentrifugation, AAVs were extracted by puncture and further concentrated using a 50 ml Amicon column (Millipore Sigma) and titrated directly on primary human T cells.

### HLA allele cross-reactivity assay

HLA allele cross-reactivity of the A2-CAR-expressing Tregs was determined based on a previously reported method (26). In brief, 2.5 × 10^4^ FACS-purified A2-CAR Tregs, as well as 2.5 × 10^4^ control untransduced polyclonal Tregs, were incubated with 0.5 μl to 5 μl PE-labeled FlowPRA Single HLA Antigen bead panels (FL1HD01 and FL1HD02, OneLambda, Los Angeles, CA), a fixable viability dye (Ghost Dye BV510, Tonbo Biosciences, San Diego, CA), and anti-CD45 e450 (clone HI30, eBioscience, San Diego, CA) for 30 minutes at 37° C. After incubation, the suspensions were washed with DPBS, fixed with 0.5% neutral buffered formalin (VWR International, West Chester, PA), washed again with DPBS, and run in a BD LSRII flow cytometer. Single antigen beads decorated with different HLAs fluoresce in the PE channel with distinct intensity, allowing one to discern the individual HLA alleles. The abundance of unbound beads was quantified in the presence of either A2-CAR Tregs or untransduced Tregs for each single HLA antigen group. Percentage relative binding of A2-CAR Tregs to each HLA allele was then calculated using the following formula

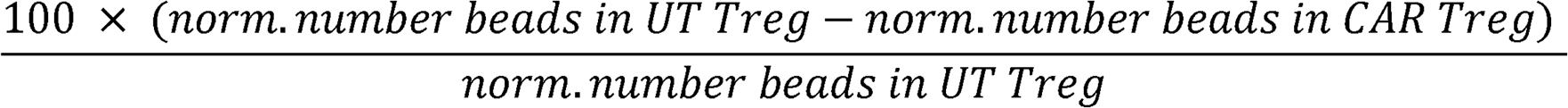

i.e. by dividing the normalized (norm.) number of beads in the untransduced (UT) Treg condition for a specific HLA minus the normalized number of beads in the A2-CAR Treg condition for that same HLA by the normalized number of beads in the untransduced Treg condition, multiplied by 100. HLA antigen bead numbers were normalized using the following formula

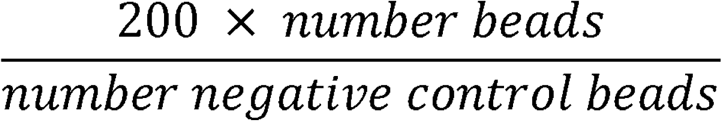

i.e. by multiplying the number of beads of interest in each HLA peak by 200, divided by the number of negative control beads in the sample, to correct for variations in the absolute number of negative control beads acquired in each sample.

### Genome engineering

CRISPR/Cas9 genome editing in Tregs and bulk T cells was carried out using ribonucleoprotein (RNP) electroporation as previously described (27). Briefly, RNPs were produced by complexing a two-component guide RNA (gRNA) to Cas9. crRNAs and tracrRNAs were chemically synthesized (Dharmacon, IDT) and Cas9-NLS (nuclear localization signal) was recombinantly produced and purified (QB3 Macrolab). Lyophilized RNA was resuspended at a concentration of 160 μM, and stored in single use aliquots at −80°C. crRNA and tracrRNA aliquots were thawed, mixed 1:1 by volume, and annealed at 37□°C for 30 min. 40 μM recombinant Cas9 was mixed 1:1 by volume with the 80 μM gRNA (2:1 gRNA to Cas9 molar ratio) at 37□°C for 15 min to form an RNP complex at 20 μM. RNPs were electroporated immediately after complexing into Tregs and T cells resuspended in supplemented P3 buffer (Lonza).

Guide RNA sequences used for gene editing were:

*TRAC*: CAGGGTTCTGGATATCTGT
*TRBC*: CCCACCAGCTCAGCTCCACG
*HLA-A2*: CCTCGTCCTGCTACTCTCGG

Following electroporation, Tregs and T cells were replated for expansion together with the AAV6 containing the A2-CAR homology-directed repair (HDR) template. Alternatively, two days after activation, Tregs were transduced with a lentivirus at a multiplicity of infection (MOI) of 1 by spinoculation for 30 min at 1200 G. The next day, the cells were washed to remove residual virus from the medium and further expanded with recombinant human IL-2 (300IU/ml). In some instances, A2-CAR^+^ cells were FACS-purified on Day 9 based on MYC-tag expression and the TCR was deleted by electroporating a CRISPR/Cas9 RNP complex targeting the constant region of the TCR beta chain (*TRBC*).

### A2-CAR Treg trafficking to transplanted pancreatic islets

Female or male NSG mice were rendered diabetic by a single intraperitoneal (i.p.) injection of streptozotocin (STZ) at 220mg/kg and islets were transplanted 72-96 hours later. Blood glucose levels were monitored 2-3 times per week using a glucometer (Nova Max Plus Blood glucose and Ketone Monitor, Nova Diabetes care, Billerica, MA). Only mice with blood glucose levels above 300mg/dl were used for transplantation. Pancreatic islets from NSG.HLA-A2 transgenic mice (A2-NSG, NOD.Cg-Prkdc^scid^ Il2rg^tm1Wjl^ Tg(HLA-A/H2-D/B2M)1Dvs/SzJ, Jackson Laboratories, Bar Harbor, ME, Stock number 014570) were isolated as previously described (28). Human pancreata were procured from deceased multi-organ donors with research use consents and approval from UCSF institutional review board. Human research islets were isolated by the UCSF Diabetes Center Islet Core following standard protocols (29). A total of either 500 mouse islets or 3000 human islet equivalents (IEQs) were transplanted under the kidney capsule or into the spleen. Blood glucose levels of < 200mg/dl on two consecutive days were defined as successful islet engraftment. Mice that only attained partial graft function (blood glucose range 200-500mg/dl) by 10 to 14 days after transplant were given subcutaneous insulin pellets (Linbit, LinShin Canada) to support graft function. Luciferase-expressing A2-CAR Tregs or A2-CAR T cells were infused intravenously in STZ-induced diabetic mice transplanted with mouse HLA-A2^+^ islets. Luciferase activity was monitored 2-3 times per week. These animals were anesthetized in an isofluorane chamber, injected i.p. with 100 μl of 15 mg/ml D-Luciferin (Biosynth, Staad, Switzerland) and, 7 min later, imaged in a Xenogen IVIS Spectrum Imaging System (PerkinElmer, Richmond, California). Luciferase data analysis was performed using Living Image software (PerkinElmer).

### Xenogeneic graft-vs-host disease (GvHD)

NOD.Cg-Prkdc^scid^ Il2rg^tm1Wjl^/SzJ (NSG) and NOD.Cg Prkdc^scid^ Il2rg^tm1Wjl^/ Tg(HLA-DRB1)31Dmz/SzJ/H2-Ab1tm1Gru x NOD.Cg-Tg(HLA-A/H2-D/B2M)1Dvs/SzJ (A2-NSG) were obtained from Jackson Laboratories. For GvHD induction, animals were irradiated (2.5Gy) 24 hours prior to retroorbital intravenous (i.v.) infusion of 5 × 10^6^ freshly isolated PBMCs from either an HLA-A2-positive or an HLA-A2-negative donor with or without 2.5 × 10^6^ *ex vivo* expanded third-party A2-CAR Tregs. All mouse experiments were performed according to a UCSF Institutional Animal Care and Use Committee (IACUC) approved protocol.

## Results

### Development of an HLA-A2-specific CAR

To engineer an anti-A2 CAR, we first cloned the variable regions of the heavy (V_H_) and light (V_L_) chains of an A2-specific IgG1κ antibody from a hybridoma (SN607D8) produced using B cells isolated from a previously described sensitized donor (16). This antibody was reported to bind to HLA serotypes A2 and A28, which includes HLA-A68 and A69 alleles. After cloning the SN607D8 scFv from the hybridoma, we evaluated phage-displayed SN607D8 scFv binding to two human tumor cell lines. The THP-1 monocytic cell line expresses HLA-A2, but not A28, whereas the RPMI 8226 myeloma cell line is HLA-A2^−^ but has a genotype of HLA-A*6802 and is thus HLA-A28^+^. The results showed that the SN607D8 scFv indeed binds to both cell lines (Supplementary Figure 1), demonstrating the retention of the original specificity of the antibody.

We then cloned the SN607D8 scFv into a construct that contained an IgG4 hinge, the CD28 transmembrane domain, and a signaling domain composed of the CD28 and CD3ζ intracellular domains. Unexpectedly, the CAR failed to express on the surface of human T cells (data not shown). To rescue the expression, we grafted the complementarity-determining regions (CDRs) of the heavy and light chains of the SN607D8 scFv into the framework regions of an scFv derived from the anti-HER2 antibody Herceptin (trastuzumab), which is known to be compatible with CAR surface expression (30). The resulting grafted heavy and light chains (Figure 1A) were connected via a 15 amino acid linker (GGGGS)_3_ to form a new grafted scFv, termed QT007YL. Automated computer modeling with an antibody structure prediction tool, ABodyBuilder (31), showed that the grafted scFv folds as expected (Figure 1B). We then generated a new A2-CAR for expression in human T cells by fusing the QT007YL scFv to an IgG4 hinge, the CD28 transmembrane domain, and a CD28-ζ signaling domain. The resulting A2-CAR was cloned with an N-terminal MYC-tag into a pCDH lentiviral vector behind an EF1α promoter (22). A truncated EGFR (EGFRt) was cloned in-frame behind the CAR separated by a 2A self-cleaving peptide to enable facile evaluation of lentiviral transduction using expression of EGFRt. To enable *in vivo* tracking of CAR-expressing cells, we also generated a version of the lentiviral construct with a luciferase gene behind the CAR separated by a 2A peptide (Supplementary Figure 2A). We first transduced HLA-A2-negative Jurkat T cells to assess the expression and function of the grafted A2-CAR. Detection of the MYC-tag via flow cytometry verified efficient surface expression of the A2-CAR (Supplementary Figure 2B). Lentiviral transduction of primary human Tregs with the grafted A2-CAR also resulted in co-detection of the MYC-tag and EGFRt on the cell surface (Figure 1C). Moreover, A2-CAR Jurkat T cells upregulated CD69 and CD25 expression specifically when co-cultured with irradiated HLA-A2^+^ K562 tumor cells (Supplementary Figure 2C), suggesting that the grafted A2-CAR is able to activate T-cell signaling in response to HLA-A2.

**Figure 1:**
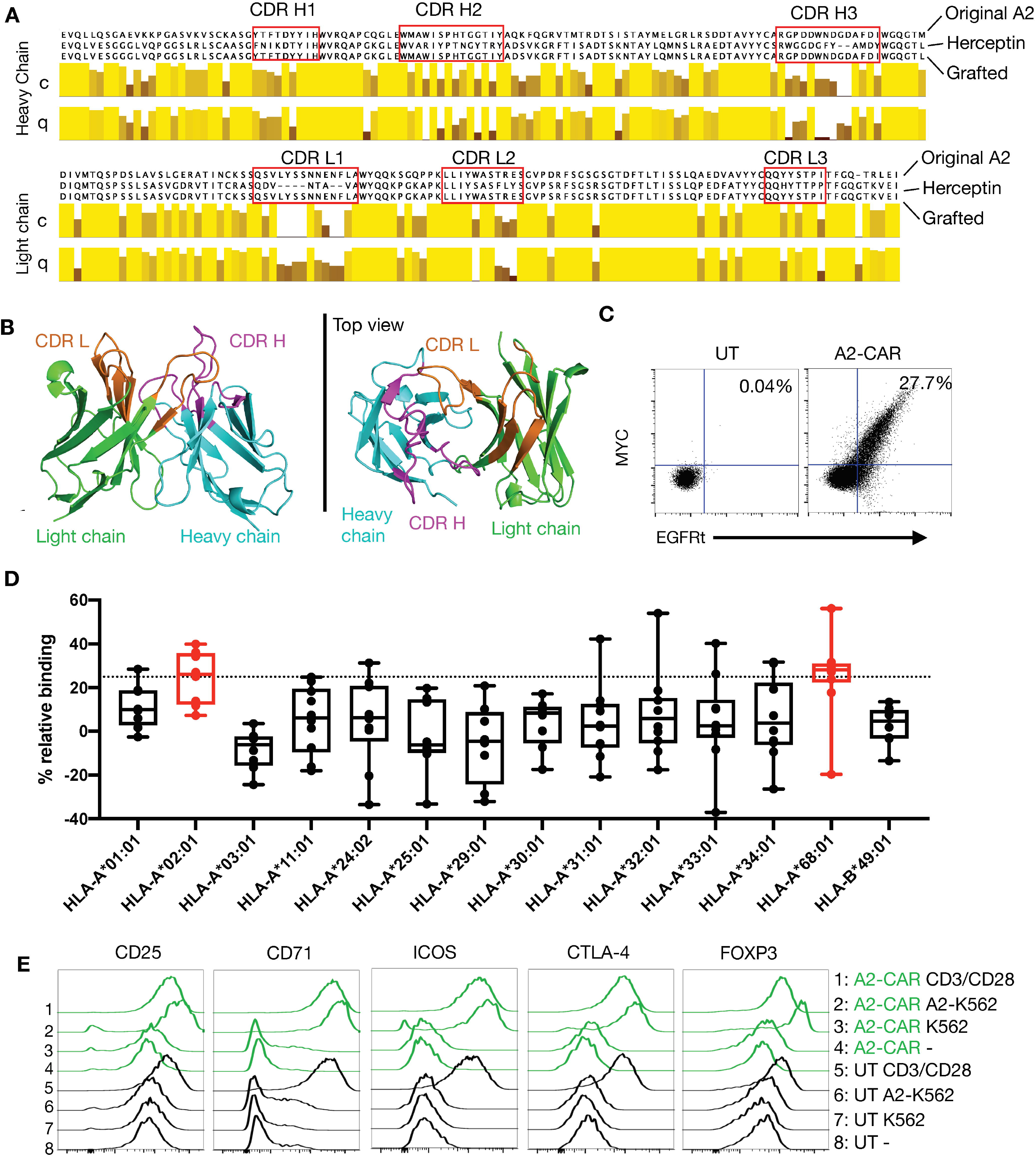
Generation of a grafted A2-CAR. **(A)** Grafting strategy comparing the original V_H_ and V_L_ chain sequences of the SN607D8 hybridoma and of the Herceptin (trastuzumab) 4D5 scaffold. The grafted amino acid sequences are shown. The sequences of SN607D8 and 4D5 HER2 were aligned using the software Jalview and the level of conservation (C) and quality (Q) of each amino acid between SN607D8 and 4D5 sequences were compared. Conservation reflects similarity of the physicochemical properties of amino acid residues. Identical residues are shown as light-yellow columns and residues with more dissimilar physicochemical properties are marked with darker column colors. Quality measures the likelihood of observing a mutation in any particular amino acid residue position (45). CDRs were predicted using Paratome (21). **(B)** The conformation of the grafted antibody was predicted with ABodyBuilder (31) and displayed using PyMOL Molecular Graphics System (DeLano Scientific, San Carlos, CA). **(C)** EGFRt and MYC-tag expression on Day 6 of culture of human Tregs transduced with the grafted A2-CAR-2A-EGFRt lentivirus. **(D)** OneLambda FlowPRA Single HLA Antigen bead panels FL1HD01 and FL1HD02. Percentage relative binding of A2-CAR Tregs to each HLA allele was calculated as described in the Materials and Methods section. Plotted averages of at least 5 independent experiments. Red coloring indicates HLA allele beads surpassing the 25% binding threshold to be considered binders. **(E)** On Day 9, A2-CAR Tregs were cultured for another 48 hours alone, with anti-CD3/CD28 beads, or with irradiated (4000 rad) parental A2^−^ K562 or A2-expressing K562 cells. CD25, CD71, ICOS, CTLA-4, and FOXP3 expression were analyzed thereafter using flow cytometry. *Abbreviations: scFv, single-chain variable fragment; CDR, complementarity-defining region; A2, HLA-A2; UT, untransduced.*

The HLA-A2 molecule contains many polymorphic eplets that are shared with other HLA class I molecules (Supplementary Table 2). The parental monoclonal antibody SN607D8 used to generate the grafted A2-CAR has a specificity for the eplet 144TKH (142T, 144K, and 145H residues), which is shared between HLA-A2, −A68, and −A69, but not with HLA-A3, −A11, or −A24 (32). To verify that the grafted A2-CAR retained the specificity for 144TKH, we tested binding of grafted A2-CAR transduced Tregs to a FlowPRA Single Antigen bead panel. In this assay, CAR specificity is defined as an increase in binding to HLA-bearing beads over control beads of at least 25%, a threshold used to define the binding specificity of a previously reported A2-CAR (26). The grafted A2-CAR reacted with HLA-A2 and HLA-A68, but not HLA-A3, −11, or −24 (Figure 1D), demonstrating that the specificity of the parental antibody was preserved in the QT007YL scFv after grafting. Finally, we evaluated A2-CAR-mediated *in vitro* activation of human Tregs following co-culture with either HLA-A2^+^ or HLA-A2^−^ target cells (Figure 1E). A2-CAR Tregs upregulated CD25, CD71, ICOS, FOXP3, and CTLA4 48 hours after stimulation with HLA-A2^+^, but not with HLA-A2^−^ target cells, further demonstrating the specificity of the QT007YL A2-CAR.

### In vivo trafficking of monospecific A2-CAR T cells in an islet transplant model

Next, we injected QT007YL A2-CAR-expressing T cells in NSG mice to determine whether CAR expression could redirect T cells to HLA-A2-expressing tissues *in vivo*. A significant fraction of human T cells can recognize mismatched HLA and trigger rejection of transplanted allogeneic human tissue in NSG mice (33). Additionally, human T cells have conspicuous reactivity against xenogeneic antigens expressed in the mouse host, with the potential to divert T cells away from human grafts and also eventually cause GvHD (34). To avoid these confounding issues, we first lentivirally transduced primary human T cells to express the A2-CAR. We subsequently generated A2-CAR^+^TCR^deficient^ T cells by CRISPR/Cas9-mediated knockout of endogenous TCR expression from Day 9 FACS-purified A2-CAR^+^ cells (Figure 2A). Because pre-existing TCRs were still expressed on the cell surface after gene deletion, we were able to expand the edited cells by restimulation with anti-CD3/CD28 beads for another 7 days, resulting in 95.65% CD3^−^ and 80.1% MYC-tag^+^ cells (Figure 2B). Co-culturing these TCR^deficient^ A2-CAR T cells with islets from HLA-A2 transgenic NSG (A2-NSG) or WT NSG mice for 48 hours resulted in the selective destruction of A2-NSG transgenic mouse islets (Figure 2C), demonstrating that the grafted A2-CAR can be specifically activated by HLA-A2 molecules expressed on islet tissue *in vitro*. To determine if TCR^deficient^ A2-CAR T cells can recognize A2-expressing islets *in vivo*, we first transplanted HLA-A2 transgenic mouse islets into STZ-induced diabetic NSG mice, and then infused the mice with 2 × 10^6^ TCR^deficient^ A2-CAR T cells after the islet grafts had been established. Kidney capsule is a standard site for islet transplantation. However, human CD4^+^ T cells efficiently trafficked to the lungs, livers and spleens, but not kidneys of NSG recipients (Supplementary Figure 3). We thus transplanted the islets either into the spleen or under the left kidney capsule and monitored luciferase-expressing A2-CAR T cell migration using bioluminescence imaging. We observed progressive increase in luciferase signal in the recipient mice, although it was difficult to discern the accumulation of A2-CAR T cells in the spleen versus the left kidney (Figure 2D). However, A2-NSG islets were rejected in both groups, with a median survival of 6 days in the spleen and 11 days under the kidney capsule (Figure 2E). This demonstrated that the A2-CAR was able to direct the trafficking and accumulation of human T cells to sites of antigen deposition outside the route of T cell migration.

**Figure 2:**
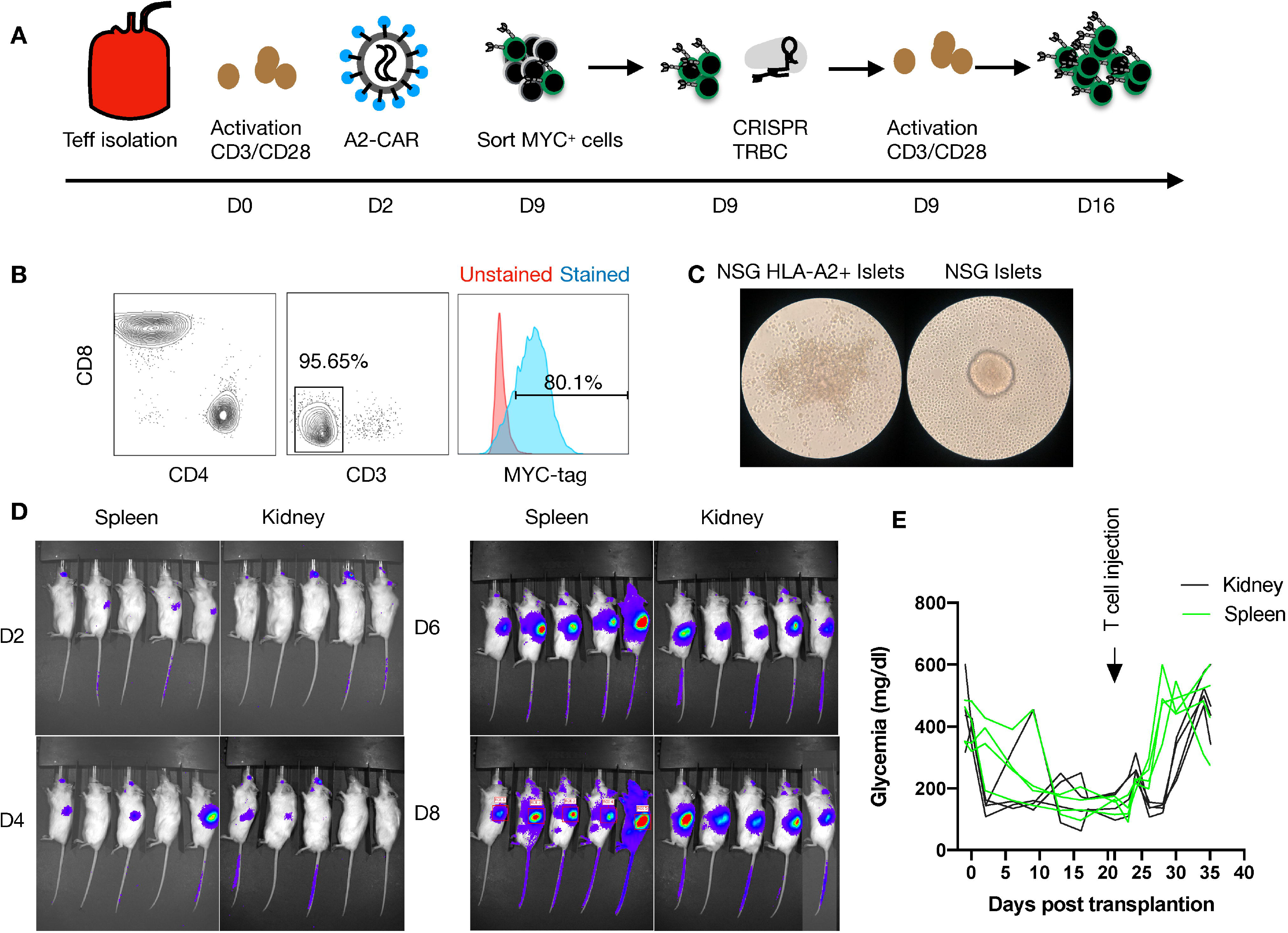
Human A2-CAR T cell trafficking and function in an islet transplantation model. **(A)** Experimental design of human T-cell engineering and expansion. **(B)** Editing efficiency as measured by CD3 and MYC-tag expression on T cells prior to infusion. **(C)** WT NSG or HLA-A2 transgenic NSG mouse islets pictured 48h after co-culture with A2-CAR T cells (100 IEQ with 1 × 10^5^ T cells). **(D)** Luciferase activity over time after infusion of 2 × 10^6^ A2-CAR T cells in mice that received an A2 transgenic murine islet transplant either under the left kidney capsule or into the spleen. **(E)** Glycemia monitoring after streptozotocin (STZ) injection, islet transplantation, and T-cell infusion (2 × 10^6^/mouse). An insulin pellet was placed subcutaneously in mice with partial graft function (blood glucose >200mg/dl) on Day 10 after transplantation.

We then used the kidney capsule islet transplantation model to evaluate the *in vivo* trafficking of A2-CAR Tregs. Luciferase-labeled human TCR^deficient^ A2-CAR Tregs were generated as described in Figure 2A and the resulting cells expressed Treg lineage markers FOXP3 and CD25 (Supplementary Figure 4). We then infused these cells into STZ-induced diabetic animals transplanted with A2-NSG transgenic islets. In this experiment, the islets were transplanted under the right kidney capsule to enable greater spatial separation from the spleen during bioluminescence imaging. To support Treg persistence in the absence of human IL-2-producing cells, mice infused with 2 × 10^6^ A2-CAR Tregs received daily i.p. injections of recombinant human IL-2 (50,000 IU/day/mouse). For comparison, we separately infused a cohort of mice with 2 × 10^6^ TCR^deficient^ A2-CAR conventional CD4^+^ T cells (Tconv), without IL-2 injection. In both groups, luciferase activity was observed first in the spleen and 3-7 days later in the right kidney (Figure 3A), demonstrating that both A2-CAR Treg and Tconv cells can traffic to the A2^+^ grafts. Importantly, islet rejection was observed in 3/4 mice that received A2-CAR CD4^+^ Tconv cells, but 0/4 mice that received A2-CAR Tregs (Figure 3B). Despite accumulation within the grafts, the lack of mouse islet destruction by the human A2-CAR Tregs suggests that CAR Tregs do not have overt toxicity against islets.

**Figure 3:**
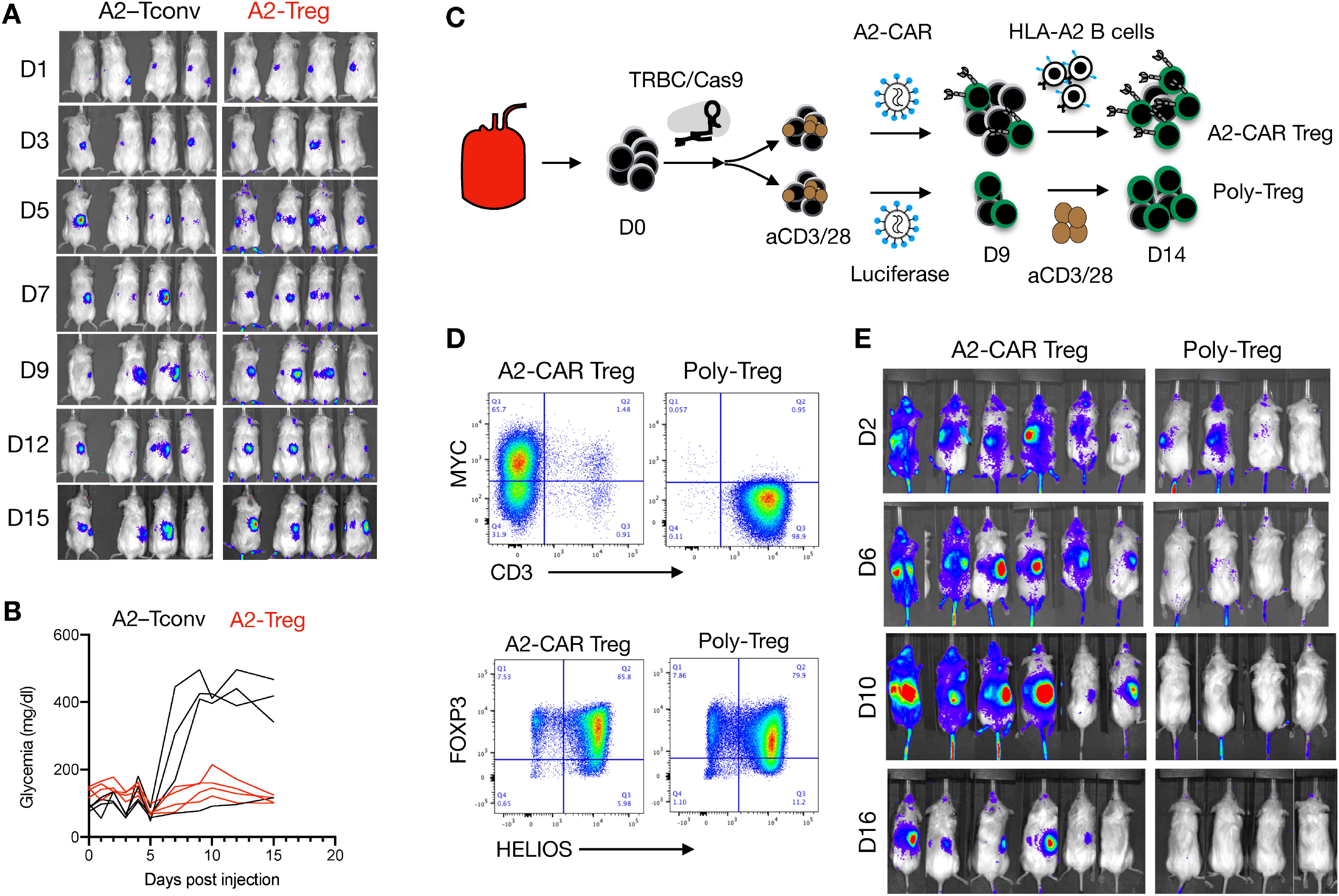
Trafficking of A2-CAR Tregs to islet grafts. 2 × 10^6^ A2-CAR CD4^+^ Tconv cells or A2-CAR Tregs (harvested on Day 10 after sorting from peripheral blood) were infused into mice that had been stably transplanted with HLA-A2 transgenic mouse islets. Luciferase activity of A2-CAR CD4^+^ Tconv cells or A2-CAR Tregs over time is shown. Glycemia was monitored over time after cell infusion. **(C)** Tregs were edited with CRISPR/Cas9 ribonucleoprotein (RNP) complexes targeting the *TRBC* locus after cell sorting, activated with anti-CD3/28 beads, and transduced with A2-CAR linked to a luciferase reporter gene by a 2A self-cleaving peptide two days later. On Day 9, Tregs were re-stimulated with HLA-A2^+^ stimulated B cells (sBCs) for another 5 days and were thereafter injected in diabetic mice (2 × 10^6^/animal) transplanted with HLA-A2^+^ human islets. In parallel, polyclonal Tregs were activated with anti-CD3/28 beads, transduced with a lentivirus expressing a luciferase reporter gene alone, and restimulated with anti-CD3/28 beads on Day 9 and injected on Day 14 in diabetic mice (2 × 10^6^/animal) transplanted with HLA-A2^+^ human islets. **(D)** Editing efficiency was measured by MYC-tag and CD3 surface expression in two independent donors. Treg purity was assessed by FOXP3 and HELIOS expression in the same donors. **(E)** Luciferase activity of A2-CAR Tregs and polyclonal Tregs (transduced with a lentivirus expressing a luciferase reporter alone) over time.

To validate that A2-CAR Tregs can traffic to human A2+ islets, we repeated this experiment with human HLA-A2^+^ islets transplanted under the right kidney capsule. In this experiment, human Tregs were first treated with CRISPR/Cas9 ribonucleoprotein (RNP) complexes targeting the TCR beta constant (*TRBC*) locus to eliminate endogenous TCR expression prior to lentiviral transduction with an A2-CAR-2A-luciferase construct. To expand sufficient numbers of TCR^deficient^ A2-CAR Tregs, cells were re-stimulated with HLA-A2^+^ stimulated B cells (sBCs) on Day 9 of culture for an additional 5 days (Figure 3C). As a control, TCR-unedited polyclonal Tregs were transduced with a luciferase-only construct and restimulated with anti-CD3/28 beads on Day 9. Tregs maintained FOXP3 and HELIOS expression prior to infusion (Figure 3D). Mice received 2 × 10^6^ Tregs intravenously with subsequent daily i.p. IL-2 injections. A2-CAR-expressing, but not polyclonal Tregs, trafficked from the spleen to the right kidney (Figure 3E). Together, these results demonstrate that TCR^deficient^A2-CAR Tregs efficiently traffic to and accumulate in human HLA-A2^+^ islets *in vivo*.

### Knocking the A2-CAR into the TRAC locus of Tregs

To achieve more uniform A2-CAR expression and investigate its function independently of the endogenous TCR, we employed homology-directed repair (HDR) to site-specifically integrate the A2-CAR into the TCR alpha constant (*TRAC*) locus, replacing expression of the endogenous TCR with expression of the A2-CAR (Figure 4A). CD4^+^CD25^high^CD127^low^ Tregs were FACS-purified and activated with anti-CD3/CD28 beads in the presence of IL-2 (300 IU/ml). Two days later, anti-CD3/CD28 beads were magnetically removed, and the cells were electroporated with Cas9-gRNA ribonucleoprotein (RNP) complexes targeting the *TRAC* and transduced with AAV6 encoding the QT007YL A2-CAR HDR template. Because this blood donor happened to be HLA-A2^+^, we also included Cas9-gRNA RNP designed to target the *HLA-A2* gene locus. *TRAC* and *HLA-A2* gene knockout efficiencies were approximately 85% and 95%, respectively. A minor (5%) population of MYC-tag^+^TCR^+^ cells was observed, likely resulting from incomplete *TRAC* inactivation and either monoallelic A2-CAR genomic integration into the other *TRAC* locus or off-target integration (Figure 4B). Importantly, while the percentage of A2-CAR-expressing MYC-tag^+^ (edited) Tregs was initially low (9%), A2-CAR^+^TCR^deficient^ Tregs preferentially expanded *in vitro*, presumably due to activation by residual HLA-A2 surface expression shortly after CRISPR/Cas9-mediated *HLA-A2* gene knockout. In the absence of further exogenous stimulation, 91% of cells were MYC-tag^+^ after 14 days (Figure 4C).

**Figure 4:**
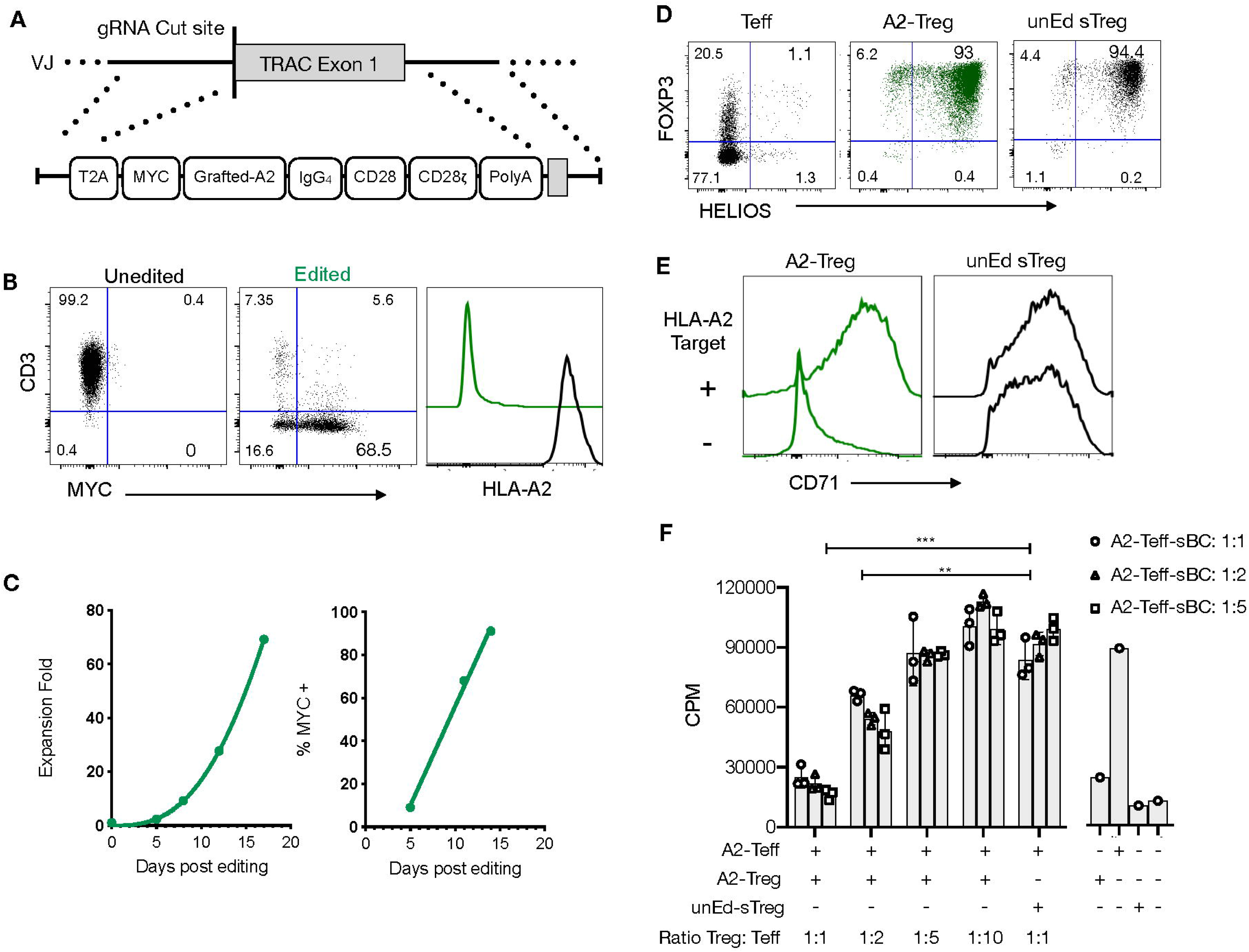
Precision engineering of an A2-CAR into the TRAC locus of human Tregs. **(A)** Homology-directed repair-mediated integration into the *TRAC* locus. The A2-CAR template was inserted using AAV6 transduction after electroporation of two CRISPR/Cas9 ribonucleoprotein (RNP) complexes targeting the *TRAC* and *HLA-A* loci. **(B)** Representative flow cytometry of the editing efficiency measured 10 days later in 3 independent experiments. CD3, MYC-tag, and HLA-A2 surface expression is shown. **(C)** Treg fold-expansion and percentage of MYC-tag^+^ Tregs over time. Fitted line plots are shown. **(D)** Fourteen days after activation, FOXP3 and HELIOS expression were assessed on edited Tregs and compared to that of unedited T cells and unedited Day 9 stimulated Tregs (unEd sTregs). **(E)** The same cells were co-cocultured with or without irradiated (4000 rad) HLA-A2^+^ NALM6 cells. CD71 expression was assessed 48 hours later. **(F)** *In vitro* suppression assays were performed using HLA-A2^+^ stimulated B cells (sBCs) as stimulator cells, A2-CAR^+^TCR^deficient^ CD4^+^ T cells as responder cells (0.05 × 10^6^ cell/96well) and A2-CAR^+^TCR^deficient^ Tregs at different ratios. After 3 days, 0.5uCi/well of ^3^[H] thymidine (Perkin Elmer, Waltham, MA) was added for the final 16 h of culture. Proliferation was assessed by ^3^[H] thymidine incorporation (counts per minute - cpm). Two-way ANOVA was used to determine the statistical significance of the difference. Data corresponds to the cells infused in Figure 5. Similar results were obtained with 3 independent donors. ** *p<0.01; *** p<0.001. Abbreviations: Teff, CD4^+^ T effector cells; A2-Treg, A2-CAR^+^TCR^deficient^ Treg; unEd sTreg, unedited Day 9 stimulated Tregs; A2-Teff, A2-CAR^+^TCR^deficient^ Teff; sBC, stimulated B cells; CPM, counts per minute.*

On Day 14 of culture, we evaluated FOXP3 and HELIOS expression among edited A2-CAR^+^TCR^deficient^ Tregs, unedited CD4^+^ Tconv, and unedited Tregs (unEd sTreg, stimulated by anti-CD3/CD28 beads on Days 0 and 9, as per our standard protocol for polyclonal Treg expansion (35)). Over 99% of A2-CAR^+^TCR^deficient^ Tregs were FOXP3 positive and 93% were HELIOS and FOXP3 double positive (Figure 4D). Co-culture of the A2-CAR^+^TCR^deficient^ Tregs with NALM6, an HLA-A2-positive B cell-derived leukemia cell line, led to a marked and specific upregulation of surface CD71 expression, demonstrating the antigen-driven activation of the edited cells (Figure 4E). Finally, we evaluated the suppressive function of the A2-CAR^+^TCR^deficient^ Tregs *in vitro* by co-culturing them with A2-CAR^+^TCR^deficient^ T cells and HLA-A2^+^ irradiated sBCs and assessing T cell proliferation. A2-CAR^+^TCR^deficient^ Tregs suppressed the proliferation of A2-CAR^+^TCR^deficient^ T cells in the presence of HLA-A2^+^ sBCs, whereas unedited polyclonal Tregs did not (Figure 4F).

### Monospecific A2-CAR Treg function in vivo in xenogeneic GvHD models

We next tested the *in vivo* functionality of A2-CAR^+^TCR^deficient^ Tregs *in vivo* within models of xenogeneic GvHD, induced by human PBMCs in sub-lethally irradiated NSG mice. In these experiments, we induced GvHD by infusing PBMCs from an HLA-A2^+^ or an HLA-A2^−^ donor into HLA-A2-transgenic or wild-type (i.e. lacking HLA-A2 expression) NSG mice (Figure 5A, Supplementary Figure 5A). This created 4 experimental groups with regards to the expression of HLA-A2: 1 absent; 2 expressed by the NSG recipients, 3 expressed by the infused PBMCs; and 4 expressed by both NSG recipients and PBMCs. A subset of mice in each of the 4 groups also received A2-CAR^+^TCR^deficient^ Tregs at the time of PBMC infusion. We first confirmed in a mixed lymphocyte reaction (MLR) that A2-CAR^+^TCR^deficient^ Tregs upregulated CD71 only in the presence of the PBMCs from the HLA-A2^+^ donor used for the immune reconstitution (Supplementary Figure 5B). To avoid direct contact between the PMBCs and Tregs during infusion, we injected the cells separately into contralateral retro-orbital plexus. A2-CAR^+^TCR^deficient^ Tregs delayed GvHD in mice that had HLA-A2 expressed by the PBMCs, the NSG recipients, or both (Figure 5B) and failed to confer any protection against GvHD in wild-type NSG animals reconstituted with HLA-A2^−^ PBMC, with a median survival of 13 days (Figure 5C). This demonstrates that A2-CAR Tregs can function in the absence of the endogenous TCR and that A2-CAR Treg-mediated protection from GvHD depends on the presence of the HLA-A2 antigen.

**Figure 5:**
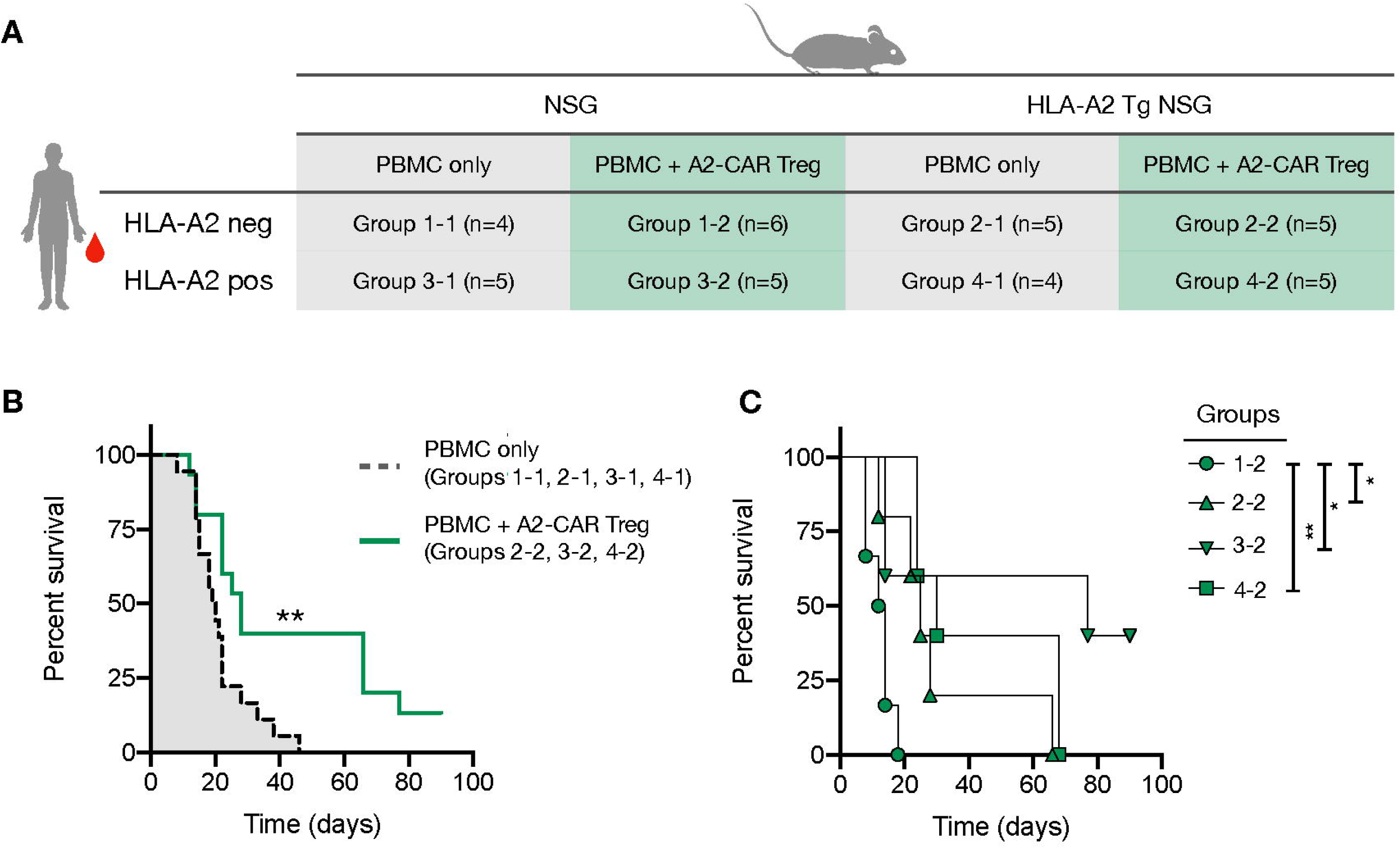
A2-CAR Tregs confer protection against graft-versus-host disease. **(A)** 2.5 × 10^6^ AAV CRISPR-edited A2-CAR^+^TCR^deficient^ Tregs were co-injected with 5 × 10^6^ A2 negative or positive PBMC in wild type or A2 transgenic immunodeficient NSG mice. PBMCs and Tregs were injected into contralateral retro-orbital plexus. **(B)** Overall survival of mice that received PBMCs (A2 positive or negative, NSG A2 positive or negative) or PBMCs and A2-CAR^+^TCR^deficient^ Tregs, excluding the condition where Tregs remained unstimulated (group 1-2). **(C)** Overall survival of mice treated with A2-CAR^+^TCR^deficient^ Tregs comparing NSG PMBC (group 1.2, round), A2-NSG PBMC (group 2-2, triangle), NSG A2^+^ PBMC (group 3-2, inverted triangle, n=6), and A2-NSG A2^+^ PBMC (group 4-2, square). Log-rank (Mantel-Cox) test was used for statistical analysis * *p<0.05,* \*\**p<0.01.*

To further investigate the mechanism of A2-CAR^+^TCR^deficient^ Treg-mediated protection, we determined the percentage of circulating human CD4^+^ and CD8^+^ T cells in the peripheral blood of treated animals at Days 7, 14, and 21 following cell injection (Figure 6A). As previously reported (10), we observed that HLA-A2^+^ T cells failed to engraft in mice that also received A2-CAR^+^TCR^deficient^ Tregs, irrespective of HLA-A2 expression by the host mice (Figure 6A-B). Meanwhile, the frequency of engrafted HLA-A2^−^ T cells was vastly reduced, but not completely eliminated, when co-injected with A2-CAR^+^TCR^deficient^ Tregs in HLA-A2^+^ transgenic host mice. In HLA-A2 transgenic mice reconstituted with HLA-A2^+^ PBMCs (groups 4-1 and 4-2), we found that the circulating HLA-A2^−^ cells, i.e. the engineered A2-CAR^+^TCR^deficient^ Tregs, remained FOXP3^+^ (Figure 6A), albeit with limitations in the number of acquired events due to the marked decrease in the number of CD4^+^ and CD4^+^HLA-A2^−^ cells over time (Figure 6B).

**Figure 6:**
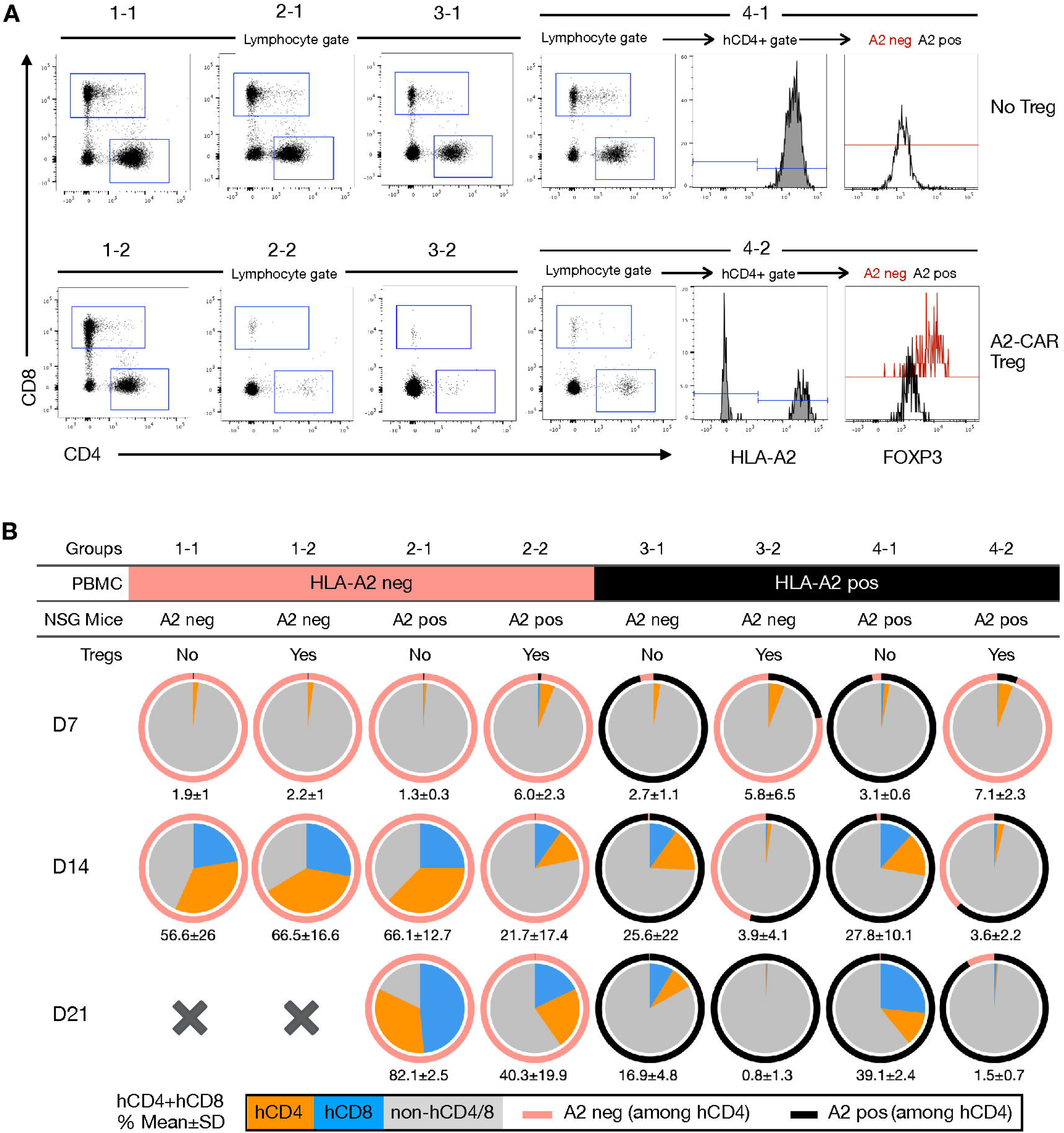
Blood monitoring of immune reconstitution in xenogeneic graft-versus-host disease. **(A)** Representative staining of human CD8^+^, human CD4^+^ and hCD4^+^A2^neg^FOXP3^+^ subpopulations after dead cell (Ghost fixable viability dye positive) exclusion. Representative example of mice from each group (1-1, 1-2, 2-1, 2-2, 3-1, 3-2, 4-1, 4-2). **(B)** Pie charts showing the mean percentage of human CD4^+^ (orange) and CD8^+^ (blue) cells in the peripheral blood of mice 7 (n=4-6), 14 (n=2-5) and 21 days (n=2-4) after injection. The numbers below each pie chart represent the mean ± standard deviation of the combined percentage of human T cells (hCD4^+^ and hCD8^+^). *Abbreviations: A2, HLA-A2*

## Discussion

Here, we report the successful development of a novel human anti-HLA-A2 CAR. Two other human and humanized A2-CARs have been previously described: one by the group of Megan Levings, where a mouse anti–A2 BB7.2 hybridoma was humanized (26), and a second by the groups of Giovanna Lombardi and Elmar Jaekel, generated from a previously published anti-HLA-A2 antibody sequence (36) (clone 3PB2 V_H_ and DPK1 V_L_) derived from a sensitized blood transfusion patient (11, 12). Our original anti-HLA-A2 hybridoma (SN607D8), first described in 2003, was isolated from a woman sensitized during her pregnancies (16). Its HLA specificity was determined by complement-dependent cytotoxicity on a large panel (n>230) of HLA-typed peripheral blood lymphocytes (37), where it was found to only cross-react with HLA-A28, a split antigen that encompasses HLA-A68 and HLA-A69 alleles. The epitope responsible for this cross-reactivity has also been pinpointed, being defined by the amino acids 142T/145H (38). Such extensive characterization was instrumental for us to confirm the preservation of the specificity of the original and the grafted A2-CAR scFv.

Our initial failure to express an A2-CAR constructed with an scFv derived from the original SN607D8 hybridoma on the surface of human T cells suggests possible conformational instability. We then grafted the scFv CDR regions into an scFv framework (trastuzumab) known to be compatible with CAR surface expression (30). Thus, the trastuzumab framework may confer greater stability to scFvs for CAR protein folding and expression. However, it should be noted that we have not tested this grafting strategy with other scFvs, and thus cannot be certain of the broader applicability of this CDR-grafting approach. Nevertheless, our success in grafting the specificity of the SN607D8 antibody shows that this approach may be useful when designing CARs for new targets.

It is currently unknown whether the endogenous TCR impacts the function of CAR Tregs. Thymically derived Tregs have a highly diverse TCR repertoire that is skewed towards recognizing autoantigens (39), and work in mice has demonstrated that Tregs require continuous TCR signaling to maintain normal immune homeostasis (40). Thus, retaining the TCR in CAR Tregs might support their homeostasis *in vivo*. However, in the context of solid organ transplantation, A2-CAR Tregs traffic to the HLA-A2-expressing graft, as shown in previous work (11, 12) and in this study, thus receiving continuous signaling via the CAR, ultimately inducing bystander suppression and supporting homeostasis independently from their endogenous TCR. Our islet transplantation experiments show that A2-CAR Tregs with or without endogenous TCR efficiently traffic to the site of antigen expression. Moreover, the finding that TCR-deficient A2-CAR^+^ Tregs suppress GvHD, in an HLA-A2-dependent manner, shows that CAR Tregs can function without their endogenous TCR.

Our results are consistent with previous work that has shown efficient GvHD prevention by A2-CAR Tregs in NSG mice infused with HLA-A2^+^ PBMCs (10). By analyzing PBMC engraftment, it was apparent that the protection from GvHD was a result of preventing T cell engraftment, possibly due to direct recognition of HLA-A2^+^ PBMCs by A2-CAR Tregs. Low T cell engraftment is not a desirable outcome of Treg therapy for GvHD, as immune reconstitution and subsequent recovery of protective immunity are needed to safeguard bone marrow transplant recipients suffering from cancer against infectious agents and residual cancer cells that escaped chemotherapy (41, 42). Our results show that A2-CAR Tregs can delay GvHD not only when the PBMCs themselves express HLA-A2, but also when HLA-A2 is expressed by the recipients and not by the PBMCs.

One potential benefit of eliminating endogenous TCR expression is to more precisely control Treg specificity, especially in the context of universal CAR Tregs for off-the-shelf use. Creating and banking such universal CAR Tregs may circumvent the challenge of expanding Tregs from immunosuppressed transplant recipients and the long production time that precludes their use in acute conditions. In this vein, we show in this study that we can simultaneously ablate HLA-A2 and TCR expression at high efficiency in human Tregs, while maintaining stable FOXP3 and HELIOS expression, as well as antigen-specific suppressive function *in vitro* and *in vivo*. Recently, progress has been made towards the generation of universal human pluripotent stem cells, which portend potential inexhaustible sources of universally compatible cells, tissues, and organs for therapy (43, 44). Our data support the feasibility of developing universal engineered Tregs with precisely controlled specificity while evading host immune destruction to induce immune tolerance. Future experiments aimed at further characterizing the suppressive potency and longevity of CAR Tregs *in vivo* will shed additional light on the efficiency, safety, and feasibility of this strategy for Treg-based cell therapy.

In conclusion, we demonstrated that it is feasible to engineer a grafted CAR directly into the *TRAC* locus of human Tregs. This strategy is highly efficient, does not cause Treg destabilization, and allowed for the generation of Tregs with CAR-restricted specificity that delayed GvHD in a target antigen-dependent manner. This strategy can be applied for precision engineering of therapeutic Tregs.

## Acknowledgements

We thank Alexander Marson for sharing the Lonza 4D 96-well electroporation system, and Juan Du, Roxxana Beltran-Valencia, and Hashim Shaikh for technical assistance.

## Author contributions

Designed the project: QT, JAB. Supervised the project: QT, JAB, JW. Designed experiments: YDM, KL, YZ. Performed experiments: YDM, LMRF, PH, GF, VN, KL, YZ, NS, AMK. Analyzed data: YDM, LMRF, QT. Provided reagents and advice: AM, FHJC. Wrote the manuscript: YDM, LMRF, QT

