## Supplementary Material for "Precision engineering of an anti-HLA-A2 chimeric antigen receptor in regulatory T cells for transplant immune tolerance"

### Supplementary Figures

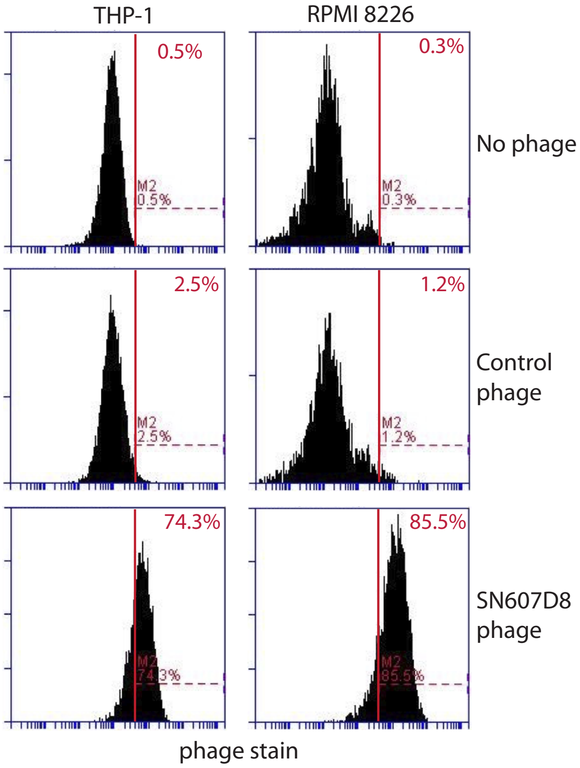


**Supplementary Figure 1: Phage-displayed SN607D8 scFv binding to HLA-A2- and HLA-A28-expressing cell lines.** The scFv from the anti-HLA-A2 hybridoma SN607D8 was cloned into a phage vector and expressed on the surface of the phage. The HLA-A2$^+$HLA-A28$^-$ THP-1 cells or the HLA-A2$^-$HLA-A28$^+$ RPMI 8226 cells were incubated with $10^7$ phages displaying the SN607D8 scFv. The cells were then stained with a biotinylated anti-phage antibody, followed by fluorochrome-conjugated streptavidin. Cell-bound phage was then analyzed using flow cytometry. Cells not incubated with phage or incubated with control phage were similarly stained and included as controls.

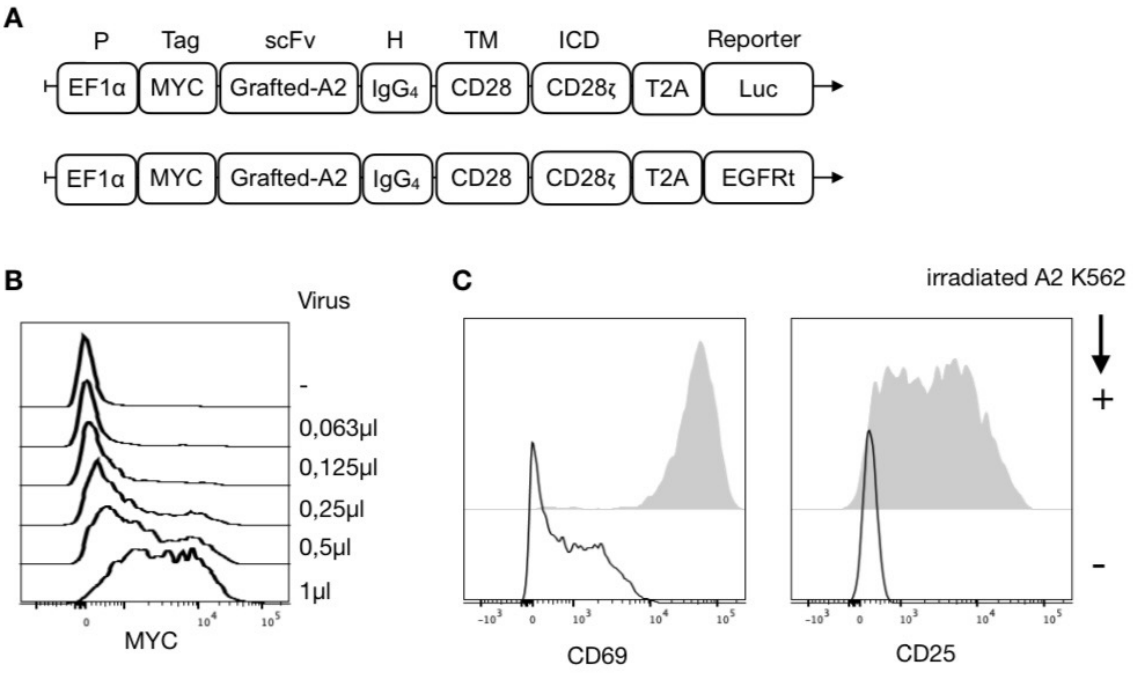


**Supplementary Figure 2: Construction of a novel A2-CAR and evaluation in Jurkat T cells. (A)** Map of the grafted anti-HLA-A2 (A2) CAR gene fused to a luciferase (luc) or an EGFRt reporter gene. **(B)** A2-CAR lentivirus was titered on 1 x $10^4$ Jurkat T cells. CAR surface expression was assessed by MYC-tag staining 72 hours post-transduction. **(C)** 2.5 x $10^4$ A2-CAR transduced Jurkat T cells were co-cultured with irradiated (4000 rad) $GFP^+$ K562 cells expressing HLA-A2 at a 1:1 ratio for 24 hours. CD69 and CD25 expression were analyzed using flow cytometry. As a control, A2-CAR Jurkat T cells were incubated alone. *Abbreviations: P, promoter; H, hinge; TM, transmembrane; ICD, intracellular domain; Luc, luciferase; A2, HLA-A2.*

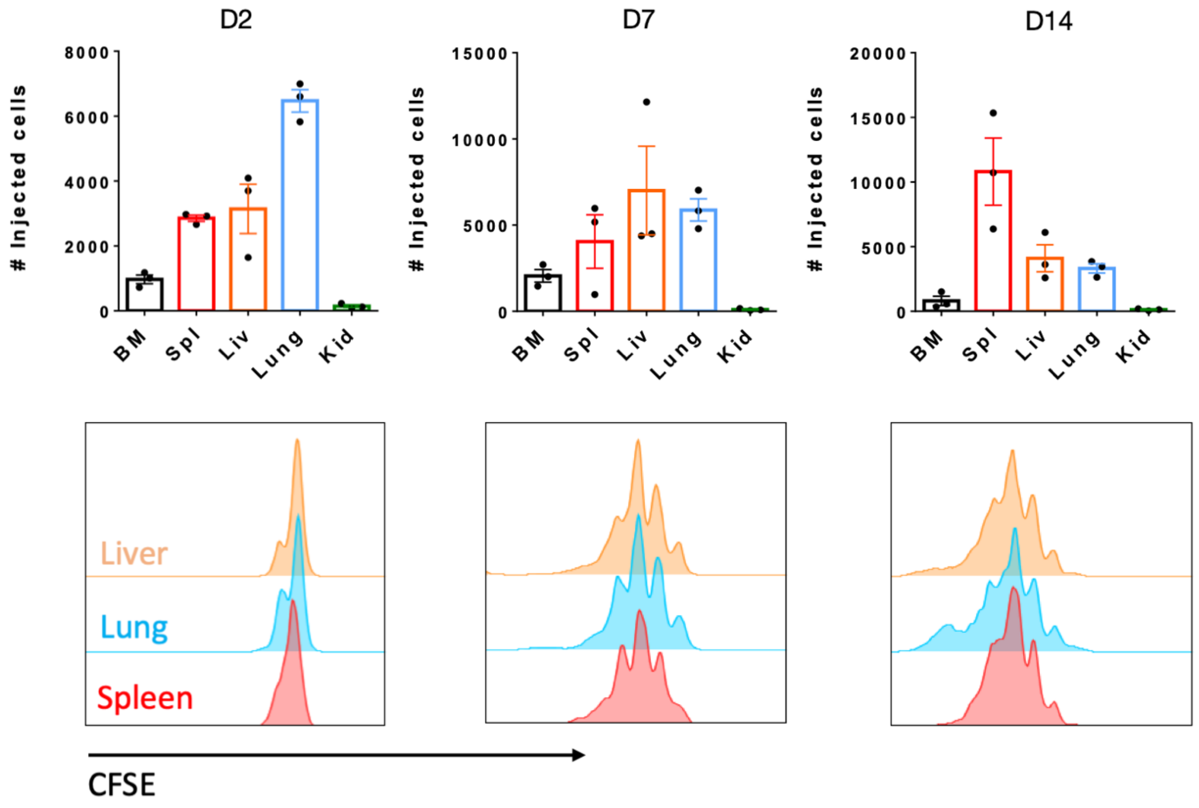


**Supplementary Figure 3: Human CD4$^+$ Tconv cell trafficking and accumulation in NSG mice**. 5 x $10^6$ CFSE-labeled human CD4$^+$ Tconv cells were injected i.v. into NSG mice. On Day 2, 7, and 14 after cell transfer, their presence (top row) and proliferation (bottom histograms) in the bone marrow (BM), spleen (Spl), liver (Liv), lungs, and kidneys (Kid) of the recipient mice were analyzed using flow cytometry.

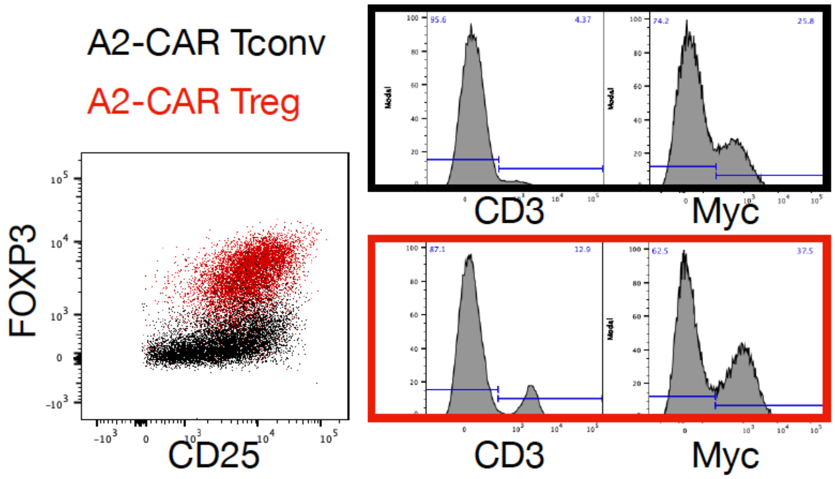


**Supplementary Figure 4: Engineering human A2-CAR$^+$TCR$^{deficient}$ Tregs.** CD4$^+$CD25$^{high}$CD127$^{low}$ Tregs and CD4$^+$CD25$^{low}$CD127$^{high}$ conventional T cells (Tconv) were sorted using FACS. Thereafter, the T-cell receptor (TCR) was deleted using CRISRPR/Cas9 and the cells were activated with anti-CD3/CD28 beads. Two days later, cells were lentivirally transduced with A2-CAR. CD25, FOXP3, CD3, and MYC-tag expression were measured on Day 9.

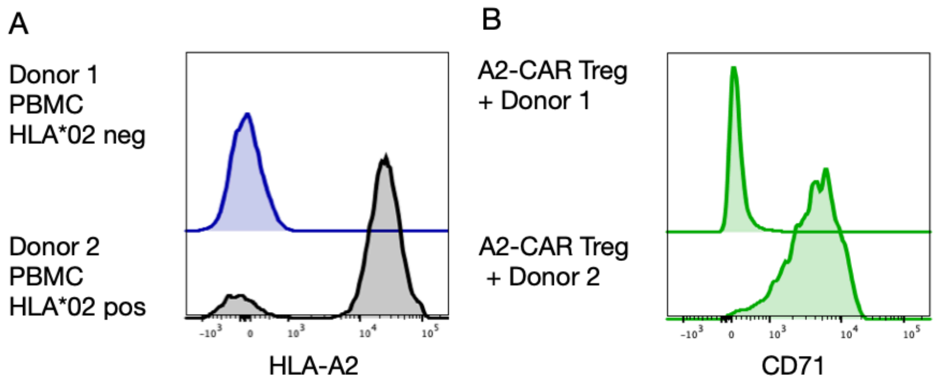


**Supplementary Figure 5: *In vitro* A2-CAR Treg activation by PBMCs from an HLA-A2$^+$ donor**. **(A)** HLA-A2 expression on PBMCs from two donors used for immune reconstitution in NSG mice in Figure 5. **(B)** A2-CAR Tregs were co-cultured for 48 hours at 1:1 ratio with the PBMCs of each donor. CD71 expression was analyzed using flow cytometry.

**Supplementary Table 1: Flow cytometry antibodies used in this study.**

| Antigen | Fluorophore | Reactivity | Clone | Company | Location |
|---|---|---|---|---|---|
| MYC-tag | A647 | Human | 9B11 | Cell Signaling Technologies | Danvers, MA |
| CD69 | PE/Cy7 | Human | FN50 | Biolegend | San Diego, CA |
| CD71 | FITC | Human | CY1G4 | Biolegend | San Diego, CA |
| CD4 | PE/Cy7 | Human | SK3 | Biolegend | San Diego, CA |
| ICOS | BV711 | Human | DX29 | BD Biosciences | San Jose, CA |
| FOXP3 | e450 | Human | PCH101 | eBioscience | San Diego, CA |
| CTLA4 | PerCP-e710 | Human | 14D3 | eBioscience | San Diego, CA |
| EGFR | PE | Human | AY13 | Biolegend | San Diego, CA |
| CD4 | PE | Human | SK3 | Biolegend | San Diego, CA |
| CD8 | PerCP | Human | SK1 | Biolegend | San Diego, CA |
| CD45 | PE/Cy7 | Mouse | 30-F11 | Biolegend | San Diego, CA |
| HLA-A2 | APC | Human | BB7.2 | Biolegend | San Diego, CA |
| HELIOS | FITC | Human | 22F6 | Biolegend | San Diego, CA |
| CD45 | e450 | Human | HI30 | eBioscience | San Diego, CA |
| Ghost | BV510 | | Viability dye | Tonbo Biosciences | San Diego, CA |

**Supplementary Table 2: Polymorphic eplets shared between HLA-A2 and other HLA class I alleles.**

| Eplet | Amino acid residues | Shared with | Reference |
|---|---|---|---|
| 44RME | 44R, 45M, 46E | A3, A11, A23, A24, A25, A26, A29, A30, A31, A32, A33, A34, A43, A66, A68, A69, A74, A80 | (16) |
| 105S | 105S | A3, A23, A24, A29, A30, A31, A33, A34, A68, A69, A80 | (16) |
| 127K | 127K | A23, A24, A68, A69 | (46) |
| 144TKH | 142T, 144K, 145H | A68, A69 | (32) |
| 145KHA | 144K, 145H, 149A | A68, A69 | (46) |
